# Design to Data for mutants of β-glucosidase B from *Paenibacillus polymyxa:* L336M, L336A, L336S, L336H, D35E, D35W

**DOI:** 10.1101/2025.05.22.655130

**Authors:** Claire Huntington, Ashley Vater, Justin B. Siegel

## Abstract

Recent groundbreaking advances in protein structure prediction have significantly propelled the fields of protein design and engineering. However, improving computational enzyme modeling remains a key area of interest, with exciting opportunities to enhance predictive capabilities. Existing algorithms face a crucial hurdle – predicting enzyme stability and function with accuracy. To address this limitation, large sets of experimental data that capture enzyme structure-function are needed to guide the training and testing of next-gen tools. The Design 2 Data (D2D) program aims to build such a data set by engaging students from around the world to contribute standardized experimental data of enzyme variants. The flagship enzyme dataset of D2D, β-glucosidase B (BglB), currently contains over 1300 records. The data consists of kinetic (*k*_cat_, K_M_, and *k*_cat_/K_M_), and thermal stability measures to expand the range of data available to train new computational algorithms. This paper explores six new single point mutants produced as part of this larger project and investigates variations in functional outcomes across the mutated sites. These data taken together with the larger and growing mutant library aim to deepen our understanding of the relationship between structure and function in BglB.

## INTRODUCTION

Computational methods are increasingly used in enzyme engineering to design novel proteins with increased activity and stability^1^. Despite recent advances in machine learning and State-of-the-art deep learning-based methods are gaining prominence for rapidly and accurately engineering novel proteins^3,4^, protein function prediction remains largely elusive^2^. Current design tools still have limited predictive ability for modeling enzyme catalytic efficiency and thermal stability (TS), and often require considerable human intuition to develop and select designs for testing ^5^.

A significant hurdle in developing computational design algorithms to enhance enzyme function and thermal stability is the lack of a comprehensive dataset with explicit measurements of catalytic parameters, (e.g., kinetic constants: *k*_cat_, K_M_, and *k*_cat_/K_M_), and protein melting temperatures (T_M_) collected in consistent reaction conditions^6^. To address this challenge, we established a national undergraduate research network to coordinate contributions to such a dataset. In this program, students capture kinetic constants and thermal stability (TS) data for single point mutations of β-glucosidase B (BglB)^7^. Program management is headquartered at University of California, Davis and has evolved to include contributions from a nationwide consortium of over forty institutions. Prior work by Carlin et al. examined the predictive abilities of several computational methods on a dataset of 100 kinetically characterized BglB single-point mutants and found weak relationships between data and the algorithms’ folding favorability scores ^6^. Two subsequent investigations involved the evaluation of 100+ designed mutants, using Rosetta and FoldX against experimentally derived stability data; Again, consistently, weak correlations between computational predictions and experimental measurements were observed^5,8^. The conclusions and recommendations of Carlin et al., and Huang et al., align: To reduce potential biases and improve the precision of functional predictions, it is essential to expand the current dataset for enzyme engineering tool improvement.

This study analyzed the kinetics and thermal stability of six new single-point mutations (D35E, D35W, L336M, L336A, L336S, and L336H) in BglB from *Paenibacillus polymyxa*. We selected two different sites to mutate to integrate structural variety into this case study, wherein D35 is part of an extensive hydrogen bond network and L336 is located in the middle of a second shell helix with the leucine side chain sitting in a hydrophobic pocket. We investigated the relationship between computational predictions of energetic changes and structural features with experimental results of catalytic and stability properties and found both expected and surprising outcomes within this small case study.

## METHODS

### BglB Mutant Design

Six BglB mutants were designed and modeled using Foldit Standalone, a graphical user interface (GUI) for foundational components of the Rosetta molecular modeling software suite^9^. Foldit Standalone employs the Rosetta energy score function for protein structure evaluation^10^. Mutations were assigned a total system energy (TSE) score and ligand energy values after using Foldit’s minimization functionality.

### Mutagenesis

A standard Kunkel method was used for site-directed mutagenesis to produce single-point BglB mutants on previously cloned pET29b+ vector following standard protocols^11^. DH5α *E. coli* transformed colonies were miniprepped and the purified plasmids were sent for Sanger sequencing (Eurofins Genomics) to verify mutagenesis success. Sequence data was analyzed with Benchling^12^.

### Protein Production and Purification

Sequence verified BglB variants were transformed, expanded, and expressed using previously described methods^8^. After induction with isopropyl β-d-1-thiogalactopyranoside (IPTG), cells were lysed chemically, and proteins were purified through nickel-based microcolumn immobilized metal ion affinity chromatography. The total protein yield was determined by measuring the absorbance at 280 nm with the A280 BioTek® Epoch spectrophotometer. Variants with protein concentrations of 0.2 mg/ml or lower were replicated to verify non-expression results. Finally, the protein purity was assessed using sodium dodecyl sulfate polyacrylamide gel electrophoresis (SDS-PAGE).

### Michaelis Menten Kinetics and Thermal Stability

The kinetic characterization of the enzyme variants involved measuring their substrate-product conversion rates using a previously established protocol where the production rate of 4-nitrophenol from the substrate p-nitrophenyl-beta-D-glucoside was monitored by absorbance at 420 nm^8^. Small variations to the Carlin et al assay method include: *p*-nitrophenyl-β-D-glucoside modified concentrations (100 mM, 25 mM, 8.33 mM, 2.78 mM, 0.93 mM, 0.31 mM, or 0.11 mM) and absorbance readings were taken every minute for fifteen minutes. Kinetic constants, k_*cat*_ and K_M_, were determined by fitting the data to the Michaelis-Menten kinetics equation using SciPy^13^.

To assess thermal stability, a fluorescence-based Protein Thermal Shift (PTS) assay was used (Protein Thermal Shift™ kit, Applied Biosystems®, Thermo Fisher Scientific) with the Quanta Studio 3 System. The purified proteins were diluted from 0.1 to 0.5 mg/mL and fluorescence was read while varying a temperature gradient from 20°C to 90°C. The resulting data were used to calculate the thermal stability parameter, T_M_, which represents the temperature at which 50% of the protein is unfolded. The T_M_ values were calculated using the two-state Boltzmann model from the Protein Thermal Shift™ Software.

## RESULTS

### Mutation Selection

Using Foldit, two sites were selected to explore mutational effects where both sites are >14Å from the active site reaction center (Figure 1); however, these sites differ considerably in overall structural compositions and neighboring intermolecular interactions.

**Figure 1.**
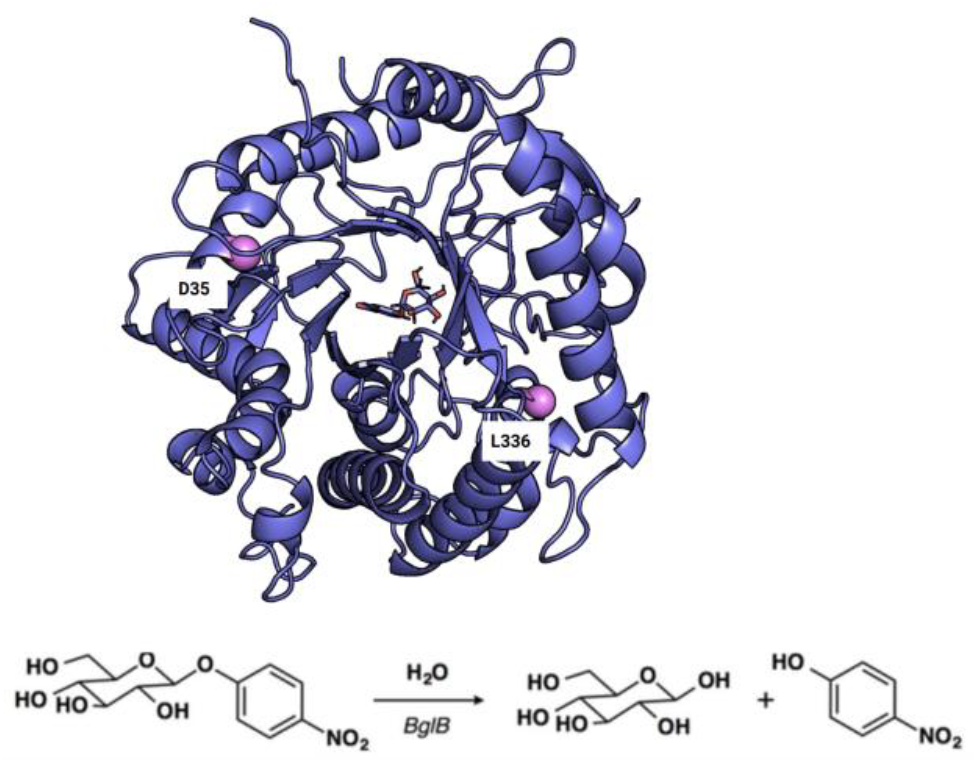
(a) Ribbon diagram of BglB (purple cartoon) with substrate 4-nitrophenol beta-D glucopyranoside (purple sticks) generated by Pymol^14^. Mutant locations D35 and L336 are labeled by pink spheres. (b) Schematic of the BglB hydrolysis reaction of reporter substrate p-nitrophenyl-beta-D-glucoside.

Site D35 is positioned 14.5 Å away from the active site and was selected due to its central role in a hydrogen bond network. Foldit modeled that this aspartic acid produced three hydrogen bonds with neighboring residues, illustrated in Figure 2A with dotted lines. We investigated two mutations at this site: glutamic acid and tryptophan. These mutants were selected to compare the effects of two very different structural moieties, particularly as they impact hydrogen bonding and if such disruptions on the protein’s surface might impact catalytic performance. Initially, the expectation was that the mutation from aspartic acid to glutamic acid would elicit negligible kinetic effects, given their structural and chemical similarities. Conversely, it was hypothesized that the introduction of tryptophan, with its marked structural unlikeness, would lead to more pronounced and deleterious effects on enzyme stability that would have a cascading effect on catalysis. Surprisingly, Foldit simulations predicted that both mutations would produce similar effects on local bonding patterns. In both cases, the mutations resulted in the loss of three hydrogen bonds with neighboring elements (Figure 2B), a phenomenon reflected in the comparable 10+ increase in their TSE scores.

**Figure 2A.**
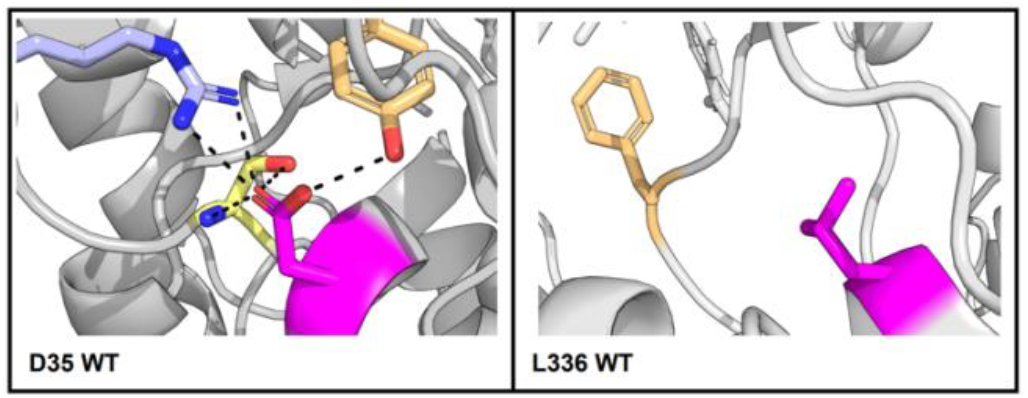
(Left-side) Pymol rendered images of site D35 and L336 wildtype highlighted in magenta. Hydrogen bonds are denoted with black dashes with neighboring amino acids for D35 with site R29 in light purple, S31 in yellow, and Y17 in tan. (Right side) L336 wildtype shown in magenta, neighbor Y294 is highlighted in light orange.

**Figure 2B.**
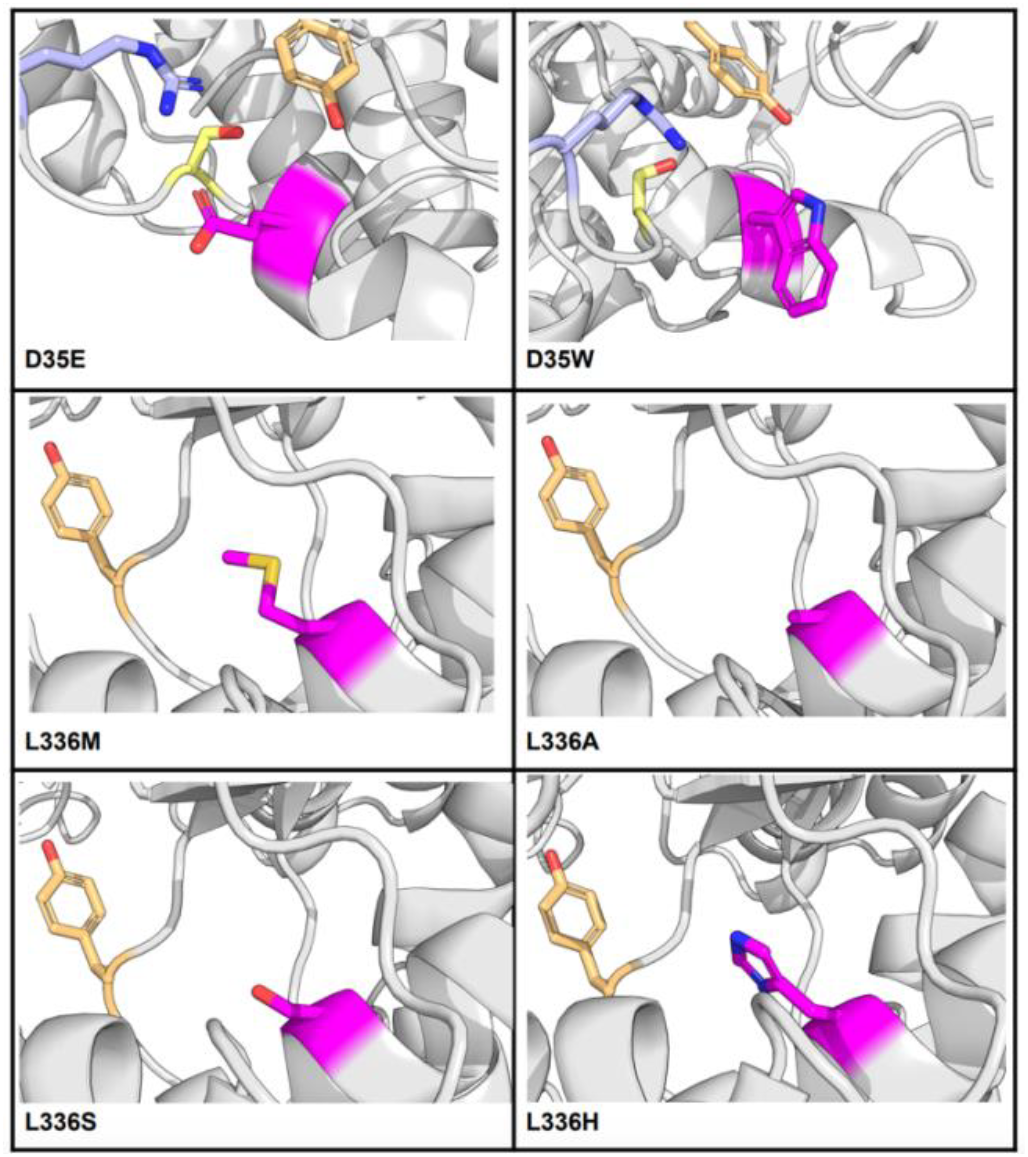
Pymol rendered images of six variant residues highlighted in pink. Residue Y294 highlighted in beige for L336 mutants. At site D35, residue R29 in light purple, S31 in yellow, and Y17 in tan.

The second mutation site L336 is situated 14.3 Å away from the active site. All mutant residues explored at position 336 were considerably structurally different from wildtype leucine. However, the native amino acid leucine, as illustrated in Figure 2A, cannot engage in hydrogen bonding with local residues and all mutations introduced at this site similarly failed to initiate any new potential hydrogen bonding interactions. Among this set of mutations, the transition from leucine to alanine represented the closest structural resemblance, while serine, despite its comparable size and structural likeness, diverges in terms of hydrophobicity. Methionine, distinguished by its sulfur-containing sidechain, appears to have no impact on total system energy. On the other hand, histidine introduces structural disruptions, resulting in an unfavorable folding state within the neighboring region, as indicated by potential atom clashing in the Foldit model with the residue Y294 (Figure 2A).

Mutations at site L336 seem to exhibit diverse effects regarding packing and clashing scores according to Foldit. Mutations to alanine and serine both reduce the packing score (−5.79 and −6.19 respectively) as compared to the wildtype (−9.55). Additionally, both these sites decrease the clashing score from the wildtype of 1.21 to 0.46 for alanine and 0.49 for serine. These score changes indicate potentially favorable space occupation as well as fewer steric hindrance conflicts. Both L336M and L336H had higher clashing scores, as well as worsening packing scores. L336H notably increases clashing (31.348), likely due to its bulky, charged side chain disrupting local structure. These findings underscore the complex interplay between local structural alterations and overall protein stability and function. In the context of TSE scores, all mutations resulted in score increases. Particularly noteworthy is the mutant L336H, which produced the second highest ΔTSE score increase among all mutations performed, with a ΔTSE of +12.

In summary, as modeled by Foldit, the site D35 had three hydrogen bonds with neighboring amino acids; alternatively, site L336 had no predicted bonding and rather had varied packing and clashing scores (Figure 2A). By introducing a diverse set of mutations at these two sites, we explore how kinetics and thermal stability are affected across varying initial hydrogen bonding presence.

### Protein Purity and Expression

All mutants expressed protein concentrations greater than 0.30 mg/ml as determined via spectrophotometer A280 analysis. Additionally, SDS PAGE was run to confirm presence and purity of the final protein products (Figure 3). All mutants besides L336A had large, visible and bold bands; L336A had a visible but smaller band. Yields ranged from 0.316 mg/ml up to the highest, L336M, at 6.6 mg/ml. The purity of all variants and wildtype appears acceptable for further, downstream functional analysis.

**Figure 3.**
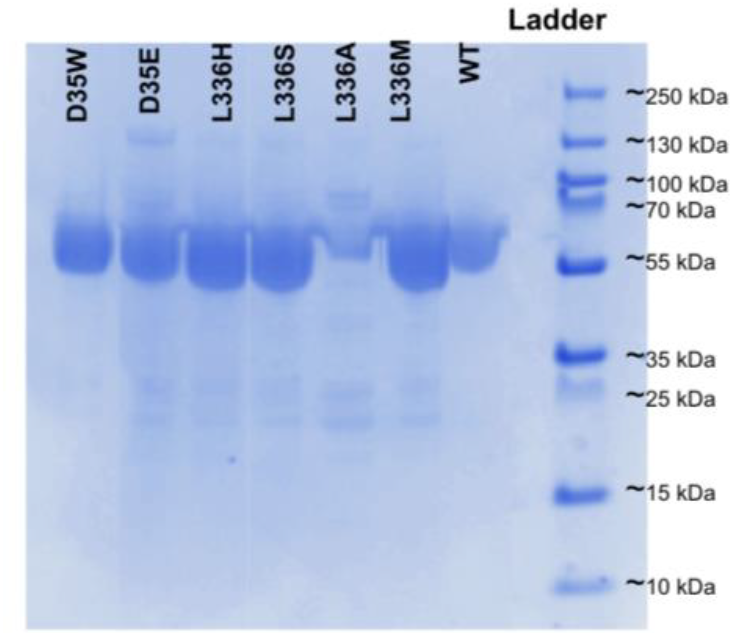
SDS-PAGE analysis of wildtype and six variant enzymes. BglB bands formed at approximately 50 kD.

### Kinetic Activity

A substrate saturation curve assay was conducted for each mutant and the wildtype protein (Figure 4). The wildtype enzyme had a *k*_cat_/K_M_ value of 218.98 min^−1^mM^−1^ which is within the normal range of values as established by previous studies^6,9^. All mutants in comparison had a lower overall kinetic efficiency ranging from 4.60 min^−1^mM^−1^ from mutant D35W to 190 min^−1^mM^−1^ with highest value from mutant L336H. All mutants in this study showed considerably reduced turnover rate, with mutations at L336 averaging only at about 60% of that of the WT and mutations at D35 reducing turnover rate by 50-fold (Table 1).

**Table 1.** Total system energy (TSE), kinetic parameters, thermal stability (T_M_), and expression (mg/mL) of six variants and wildtype.

| Protein | TSE | $k_{cat}$ (min<sup>-1</sup>) | $K_M$ (mM) | $k_{cat}/K_M$ (min<sup>-1</sup>mM<sup>-1</sup>) | $T_M$ of (°C) | Expression (mg/mL) |
| --- | --- | --- | --- | --- | --- | --- |
| L336M | -1087 | $395 \pm 18.1$ | $4.72 \pm 0.82$ | 83.7 | $47.4 \pm 0.7$ | 6.60 |
| L336A | -1086 | $568 \pm 14.9$ | $24.3 \pm 1.65$ | 23.4 | $45.2 \pm 0.5$ | 0.32 |
| L336S | -1083 | $572 \pm 20.9$ | $4.44 \pm 0.62$ | 129 | $41.8 \pm 1.0$ | 4.57 |
| L336H | -1077 | $675 \pm 13.3$ | $3.55 \pm 0.26$ | 190 | $42.3 \pm 0.6$ | 4.38 |
| D35E | -1078 | $16.3 \pm 0.40$ | $2.72 \pm 0.31$ | 5.90 | $43.1 \pm 0.8$ | 1.99 |
| D35W | -1075 | $13.0 \pm 0.60$ | $2.80 \pm 0.55$ | 4.60 | $42.1 \pm 0.9$ | 0.45 |
| WT | -1089 | $861 \pm 14.5$ | $3.93 \pm 0.26$ | 219 | $45.0 \pm 1.0$ | 5.61 |

**Figure 4.**
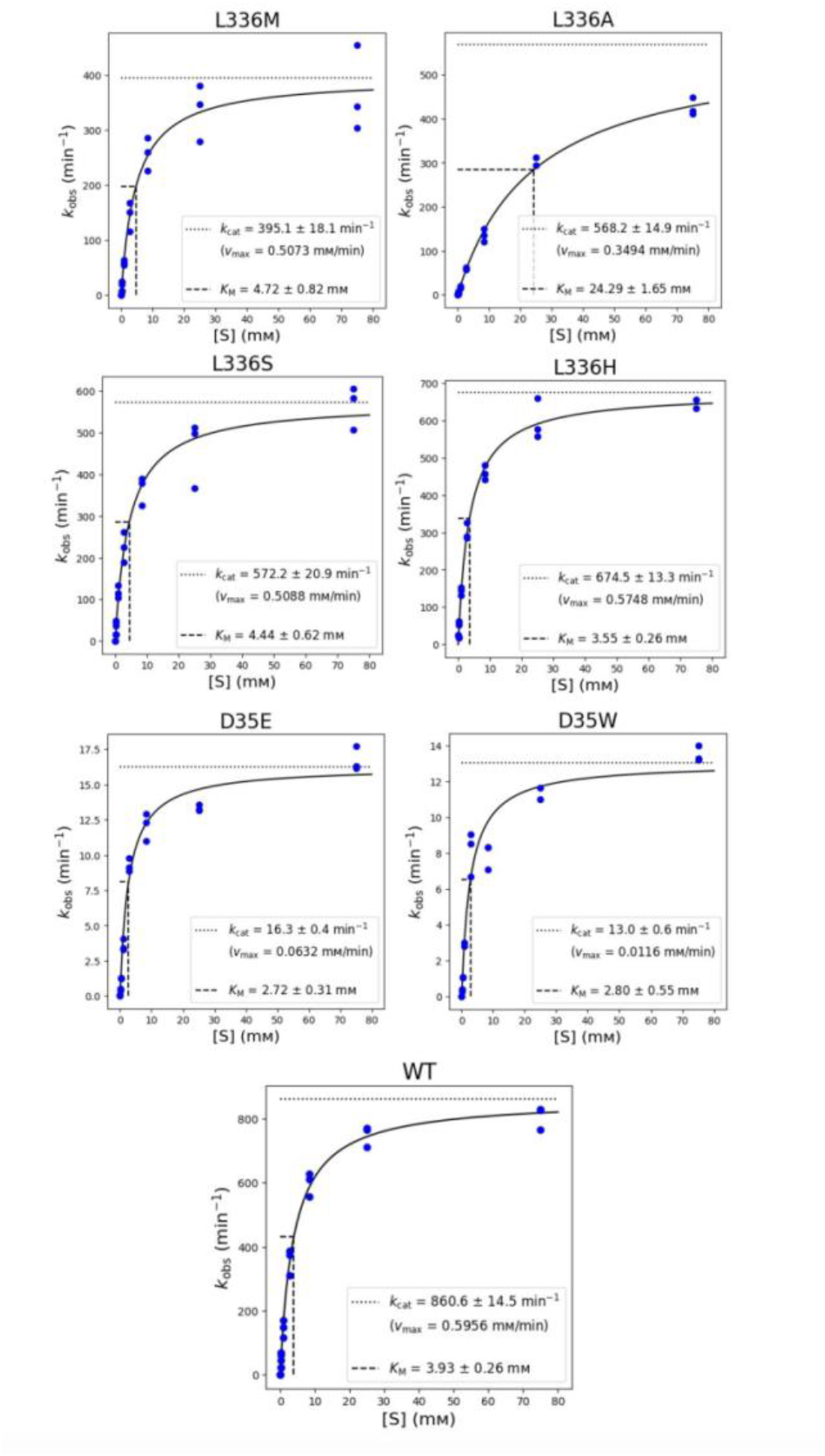
Michaelis-Menten graphs of the WT and six mutants. Concentrations of pNPG are plotted on the horizontal axis and k_obs_ on the vertical axis. The data was fitted to the Michaelis-Menten model with line of best fit for triplicate data shown.

### Thermal Stability

In order to assess thermal stability, Protein Thermal Shift assay was run in triplicate for each mutant to obtain mean T_M_ values (Figure 5). The average T_M_ value for the wild-type BglB was 44.9°C. Most mutants exhibited reduced thermal stability, with the greatest decrease observed in L336S, which had a T_M_ of 41.8°C. Only mutant L336M displayed an increased T_M_ value (+2.4°C) compared to the wildtype. The T_M_ for L336A was quite conserved with a small increase of only 0.2°C from that of the WT.

**Figure 5.**
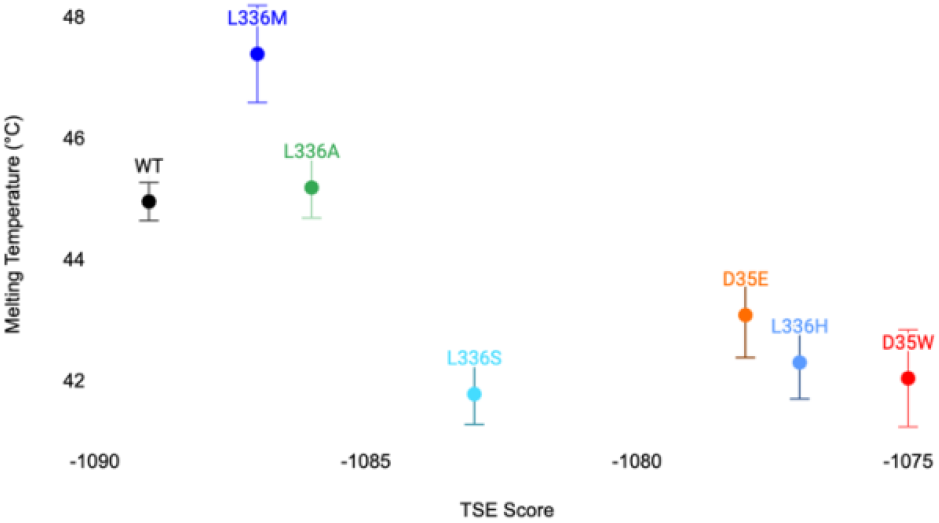
Distribution of T_M_ values in degrees Celsius for each mutant and WT plotted against the TSE score predicted by Foldit. Each mutant is represented by a distinct color and the x-axis shows degrees Celsius (°C) ranging from 42°C to 47°C. The y-axis shows TSE score ranging from −1090 to −1075.

The scatter plot depicts the relationship between T_M_ values and TSE scores for the mutants and WT. More negative TSE scores indicate enhanced predicted protein folding stability. There is a weakly observable trend: in this sample set, higher T_M_ values appear correlated with a lower---WT-like---TSE scores,. This finding, though certainly limited, suggests that protein stability, indicated by higher T_M_ values, may correlate with lower and more favorable TSE scores. Specifically, mutants such as L336M, L336A, and L336S exhibit higher T_M_ values and lower TSE scores, indicating that these substitutions likely minimally disrupt the native structure.

## Discussion

This study aimed to investigate the catalytic efficiency and thermal stability of six designed variants of β-Glucosidase B in two distinct regions of the protein, represented by site D35 and L336. These were selected to assess how changing hydrogen bonding patterns distal from the active site affect any activity and stability properties.

### Protein purity and expression

All mutants in this study expressed soluble protein, yielding protein concentrations ranging from 0.32 mg/ml to 6.60 mg/ml. By disrupting the hydrogen bonds of D35 an observable, clear reduction in expression was expected to occur due to likely destabilization of the folded state. Further, previous results from mutants at sites central to large hydrogen bond networks (e.g. D63) networks have yielded undetectable soluble protein^5^. In this case while there was clear reduction in expression for the majority of the mutants, the protein structure remained stable enough to fold and maintain activity. The only mutant that did not result in a marked reduction in expression was L336M, which is consistent with the similar molecular structure and polarity of leucine and methionine.

### Kinetics

Notable findings from kinetic data emerged across the set of six mutants, particularly from the variant’s turnover rate (*k*_cat_).

At site D35, the wildtype aspartate participates in three hydrogen bonds, both mutations at this site were predicted by Foldit to disrupt two of these bonds. For both the mutants, *k*_cat_ was strikingly reduced by 60-fold when compared to the wildtype. It is interesting that both mutants at this site had such marked decreases because while we might be expect the more dramatic sidechain moiety change from aspartic acid to tryptophan to impact the structural integrity, the change from aspartic acid to glutamic acid produced nearly identical results. Notably, the aspartate (D35) is predicted to form a hydrogen bond with its neighbor R29 (light purple in Figure 2A) which is buried deeper into the protein structure. The substitution of glutamic acid in this interaction could impede the normal folding of the protein core, potentially affecting its turnover rate capacity.

In contrast to site D35, mutants of L336 were expected to have relatively negligible effects on function, given more modest changes in TSE and no observable changes in hydrogen bonding. Indeed, the largest effect on turnover rate in this set was 2-fold less than the wildtype—versus the 60-fold of the D35 mutants.

Among the six mutants analyzed in this study, L336H was intriguing as functional results were counterintuitive to the changes in thermodynamic stability predicted by the Foldit. L336H scored unfavorably in Foldit (ΔTSE of +12). This score increase was largely attributed to the clashing of neighboring amino acid Y294 (Figure 2A) and in Foldit, attempts to resolve the clash via relaxation led to unfavorable backbone strain. Due to hydrophobicity of the region around site L336, we hypothesized that adding a polar residue like histidine would disrupt normal protein folding and thereby affect the measurable functional outcomes. However, L336H had the most conserved *k*_cat_ and K_M_ of the five mutants in this study. These data, in this particular case, inexplicably fail to support our hypothesis on the functional outcomes of this mutant.

Across the six mutants included in this study, most of the K_M_ results were unremarkable with one exception: L336A, showed a substantial increase in K_M_ (24.3 mM), a 6-fold change from the WT, but the *k*_cat_ only decreased by about 40%. It appears that the substitution of leucine with the smaller amino acid alanine at this position markedly disrupted the binding affinity of the enzyme. This was the second surprising result and was similarly challenging to rationalize given the structural and chemical parallels between leucine and alanine and the distance from the active site. Further, L336A produced virtually no score change in Foldit, which might have alerted us to a potential structural issue.

### Thermal Stability

Overall, most of the mutants in this set reduced the thermal stability of the enzyme. However, two mutants seemed to protect and even increase thermal stability of the enzyme. Mutants L336M and L336A resulted in T_M_ change of Δ+2.4 and Δ+0.2, respectively. It follows that, methionine and alanine, which share the same hydrophobicity of wildtype leucine, seem to preserve stability. As noted, when compared to the Foldit energetics there appears to be the expected relationship between change in TSE and change in T_M_, albeit more data would be needed to make statistically meaningful interpretations about this relationship.

## Conclusion

In summary, the investigation of the six mutants in this study has incrementally enriched the dataset capturing high resolution BglB structure-function relationships. Notably, mutations introduced at site D35, characterized by multiple hydrogen bonding sites, exhibited more deleterious effects on enzyme kinetics compared to mutations at site L336, wherein we could explore the effects of changes to packing and clashing. These findings underscore the crucial role of hydrogen bonding not only in enzyme-substrate interactions but also in the structural arrangement of the enzyme, emphasizing the value of scrutinizing individual positions within a protein for insights into enzyme functionality. Additionally, while four of the six variants’ data supported our functional outcome hypothesis based on TSE score changes generated by Foldit, two of the six did not. This discrepancy underscores the need for enhancing the precision of computational scoring systems, emphasizing the importance of collecting extensive datasets. The insights gained from these mutant variants contribute to ongoing efforts to catalog the structural effects on enzyme function and advance predictive software for protein modeling.

## Acknowledgments

This work was supported by the University of California Davis, the National Science Foundation (award nos. # 2315767, 2118138 and 1827246). The content is solely the responsibility of the authors and does not necessarily represent the official views of the National Science Foundation, or UC Davis. Further, this work references and is contextualized by the Design to Data (D2D) dataset, which has been built through contributions of over 1000 undergraduate students over the past decade.

## Notes

### Competing Interest Statement

The authors have declared no competing interest.

